# EXTENDING THE UTILITY OF THE WHO RECOMMENDED ASSAY FOR DIRECT DETECTION OF ENTEROVIRUSES FROM CLINICAL SPECIMEN FOR RESOLVING POLIOVIRUS CO-INFECTION

**DOI:** 10.1101/097857

**Authors:** Temitope Oluwasegun Cephas Faleye, Moses Olubusuyi Adewumi, Naomi Princess Ozegbe, Oluwaseun Elijah Ogunsakin, Grace Ariyo, Faith Wuraola Adeshina, Oluwaseun Sarah Ogunga, Similoluwa Deborah Oluwadare, Johnson Adekunle Adeniji

## Abstract

In a polio-free world there might be reduced funding for poliovirus surveillance. There is therefore the need to ensure that enterovirologist globally, especially those outside the global polio laboratory network (GPLN), can participate in poliovirus surveillance without neglecting their enterovirus type of interest. To accomplish this, assays are needed that allow such active participation.

In this light, we used 15 previously identified enterovirus isolates as reference samples for assay development. The first eight were enterovirus species B (EV-B). The remaining seven were EV-Cs; three of which were poliovirus (PV) 1, 2 and 3, respectively. A 16th sample was compounded; a mixture of two EV-Bs, three PVs and one nonPV EV-Cs (all part of the 15). In all, four samples contained PVs with the 16th consisting of mixture of the three PV types. All were subjected to the WHO recommended RT-snPCR assay, and five other modified (with substitution of the second round PCR forward primer) assays. The new primers included the previously described Species Resolution Primers (SRPs; 187and 189) and the Poliovirus Resolution Primers (PRPs: Sab 1, 2 and 3). All amplicons were sequenced and isolate identity confirmed using the Enterovirus Genotyping Tool.

The PRPs detected PV types in only the four samples that contained PVs. In addition, it was able to show that the sample 16 (mixture) contained all the three PV types. On the other hand, though the SRPs and the WHO assay also detected the three singleton PVs, in sample 16, they both detected only one of the three PV types present.

This study describes a sensitive and specific utility extension of the recently recommended WHO RT-snPCR assay that enables independent detection of the three poliovirus types especially in cases of co-infection. More importantly, it piggy-backs on the first round PCR product of the WHO recommended assay and consequently ensures that enterovirologists interested in nonpolio enteroviruses can continue their investigations, and contribute significantly and specifically to poliovirus surveillance, by using the excess of their first round PCR product.

## INTRODUCTION

In 1988, the World Health Assembly resolved to eradicate poliovirus (WHO, 1988), and as at today circulation of indigenous wild poliovirus has remained uninterrupted in only three countries (Afghanistan, Nigeria and Pakistan) globally (www.polioeradication.org). The Global Polio Eradication Initiative (GPEI) has been the major vehicle driving this effort using both immunization and active surveillance. The surveillance programme has been centred on finding poliovirus in the stool specimen of AFP cases and sewage contaminated water. The detection and confirmation of poliovirus is however, done globally in about 150 WHO accredited laboratories referred to as the Global Polio Laboratory Network (GPLN).

Detection and identification of polioviruses in specimen submitted to the GPLN is done by firstly isolating the virus in cell culture. Subsequently, the isolate is identified as a poliovirus using an array of assays ranging from serology using monoclonal virus specific antibodies to nucleic acid based tests as stipulated in the protocol (WHO, 2003; 2004). Ultimately, definitive identification of any isolate is done by nucleotide sequencing and phylogenetic analysis of the VP1 gene (WHO, 2003; 2004).

The strength of this cell culture based algorithm is the ability to detect poliovirus even at very low titre. However, a major draw-back is its dependence on infectious particles. This can constitute an avenue for short-changing the surveillance effort if the specimens are not appropriately preserved to ensure that infectious particles are viable on arrival in the laboratory. The impact of this might not be evident in developed economies where refrigeration is readily available. In resource-limited economies, where the reverse is the case, this is accommodated by moving specimen in ice chest from sampling site to laboratories. This system has challenges that range from getting ice for the chest to ensuring that refrigeration temperature in maintained in the chest from sampling site to the laboratory which may be up to hundreds of kilometres apart in a tropical climate like Nigeria. In such situation, poliovirus particles in specimens that were not well handled in transit might no longer be infectious on arrival in the laboratory. Consequently, the laboratory might not be able to detect poliovirus in such specimen. This (alongside the ongoing unrest in the three countries [Afghanistan, Nigeria and Pakistan] where indigenous circulation of wild poliovirus has remained uninterrupted) can partly account for the sporadic detection of poliovirus strains referred to as orphan polioviruses (poliovirus strains that have been previously deemed eliminated due of lack of detection for over one year)

In this light, the WHO recently recommended the Reverse Transcriptase–seminested Polymerase Chain Reaction (RT-snPCR) assay described by Nix and colleagues (Nix et al., 2006) for direct (cell culture independent) detection of enteroviruses from clinical specimen (WHO, 2015). The Nix et al., (2006) assay is an upgrade (seminested and Consensus Degenerate Hybrid Oligonucleotide Primers [CODEHOP]) version of the Oberste et al., (2003) assay. It has been shown that this algorithm is very sensitive for cell culture independent enterovirus detection and identification (Faleye et al., 2016a, Rahimi et al., 2009, Sadeuh-Mba et al., 2014). However, we (Faleye et al., 2016b) have recently shown that the assay lacks the capacity to resolve enterovirus types present in cases of co-infection. Significantly, the prevalence of enteroviruses co-infections in Nigeria is underscored by the independent emergence of 29 lineages of circulating Vaccine Derived Polioviruses (cVDPVs) between 2004 and 2014 (Pons-Salort et al., 2016), most of which are of recombinant origin (Burns et al., 2013). This necessitates the need for assays that can detect and resolve enterovirus co-infections.

Further, we had also shown (Faleye et al., 2016b) that by including primers 189 and 187 (subsequently referred to in this study as Species Resolution Primers) in the second round PCR of the WHO recommended RT-snPCR assay the resolving power of the assay could be improved. However, in a polio-free world, that there might be reduced funding for enterovirus surveillance, assays are needed that can in one swoop detect and resolve enterovirus co-infections, including different poliovirus types. Therefore, in this study, we extend the utility of the WHO recommended RT-snPCR assay by inclusion of three poliovirus-specific forward primers in the second round assay, thus, making it possible to independently detect the three poliovirus types even in cases of co-infection.

## METHODOLOGY

### Samples

Sixteen enterovirus isolates (Table 1) previously recovered from sewage contaminated water were used as reference samples in this study and analyzed as depicted in the study algorithm (Figure 1). Isolation and characterization of samples 1 through 12 have been previously described (Faleye and Adeniji 2015). Briefly, samples 1 through 8 are enterovirus species B (EV-B), and were isolated on RD cell line (Faleye and Adeniji 2015). Samples 9 to 12 are enterovirus species C (EV-C) and were isolated on MCF-7 cell line (Faleye and Adeniji 2015). Samples 13 to 15 are Sabin poliovirus 1-3 respectively. Samples 13 and 15 were isolated and characterized by the WHO Environmental Surveillance (ES) laboratory in Ibadan, Nigeria and provided to us as references for Sabin 1 and 3 polioviruses, respectively. Sample 14 (a poliovirus 2) was isolated as part of a previous study (Adeniji and Faleye, 2014) but identified subsequent to the study. Sample 16 is a mixture of six isolates, two species B (samples 1 and 3) and four species C (samples 11, 13, 14 and 15) (Table 1).

**Figure 1:**
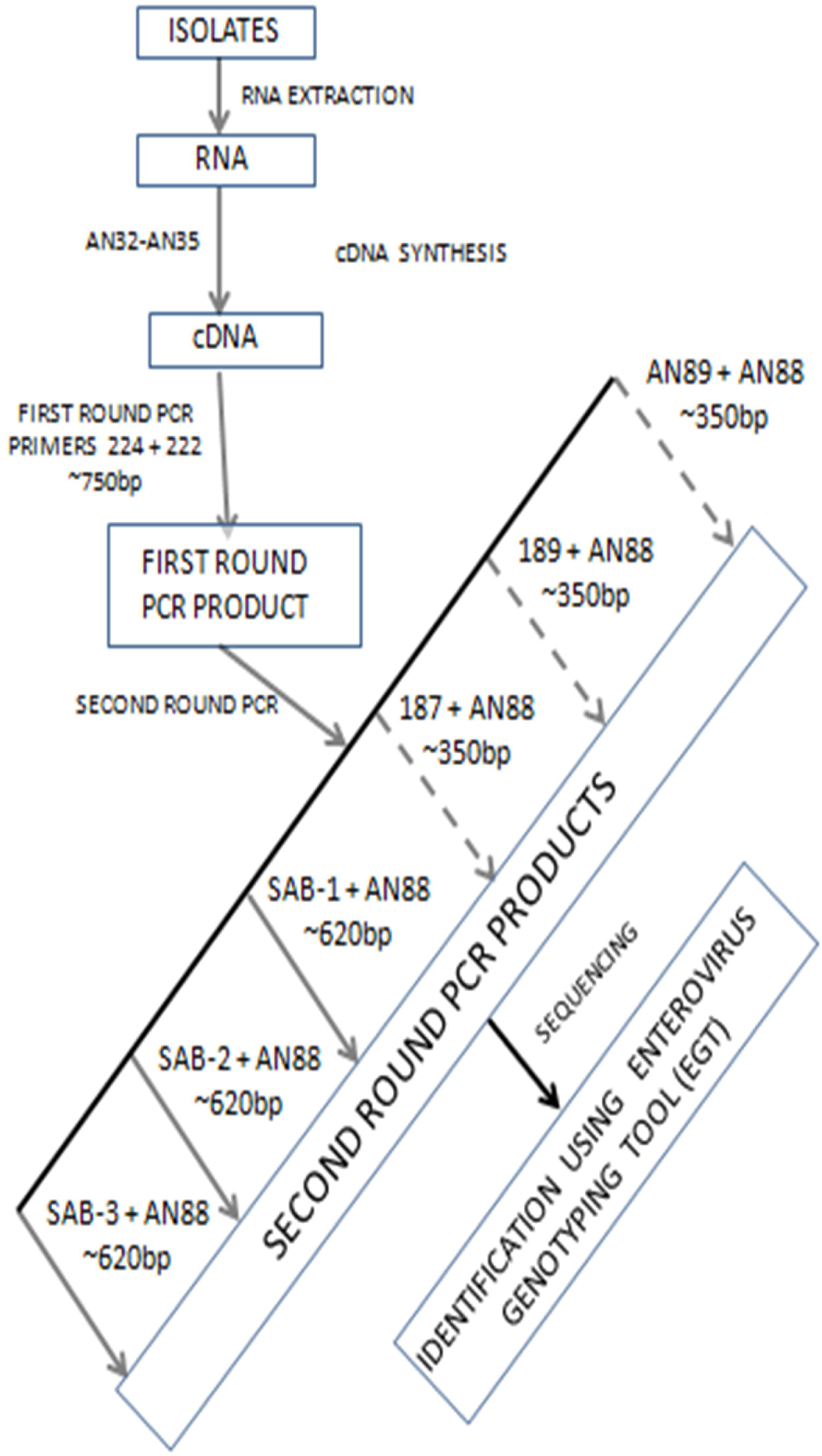
The algorithm followed in this study.

**Table 1:**
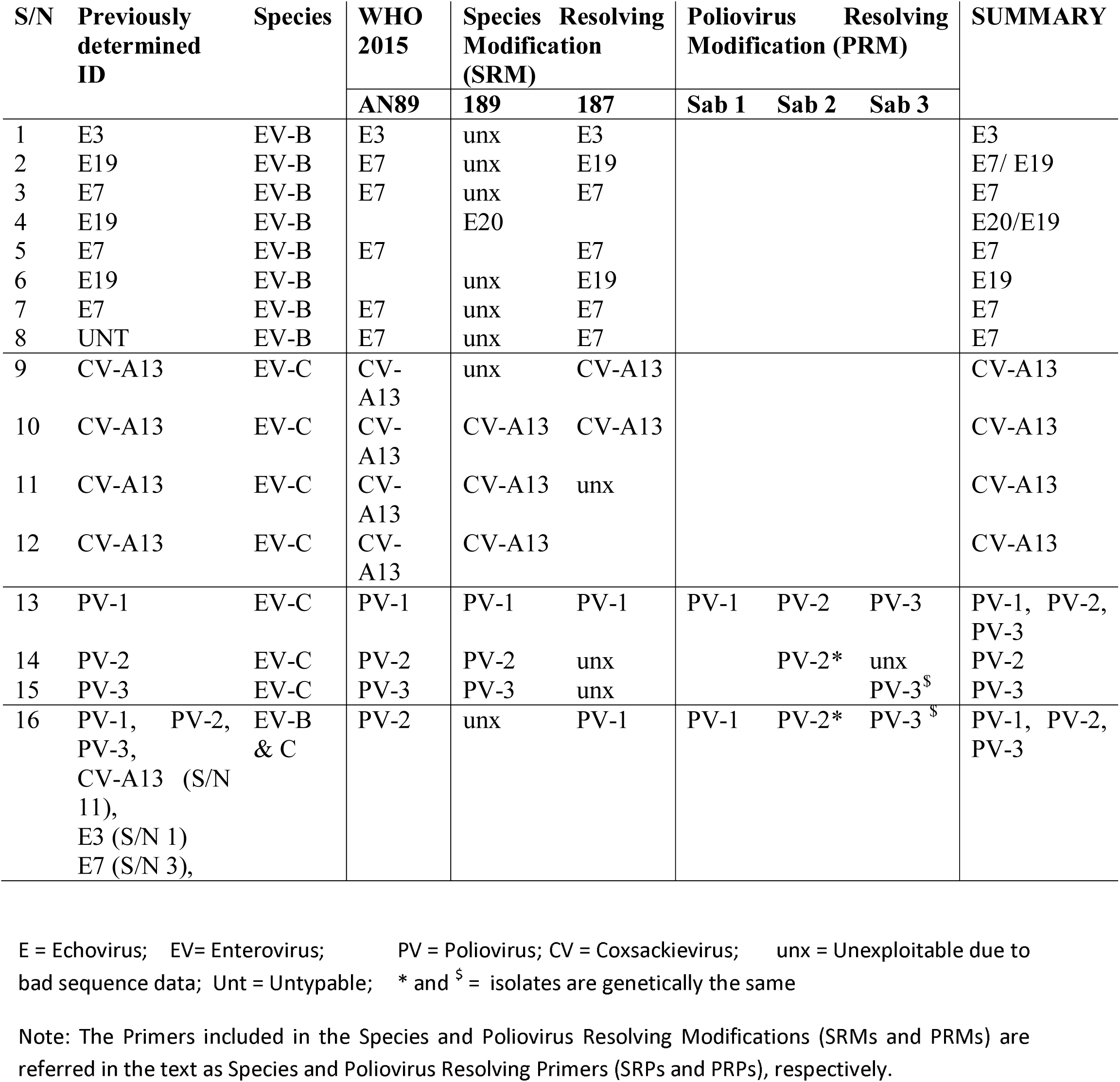
Identity of Isolates determined using the algorithm.

### RNA Extraction and cDNA synthesis

In accordance with the manufacturer’s instructions, all samples were subjected to RNA extraction and cDNA synthesis using JenaBioscience RNA extraction kit and Script cDNA synthesis kit (Jena Bioscience, Jena, Germany) respectively. Primers AN32, AN33, AN34 and AN35 (Nix et al., 2006) were used in combination for cDNA synthesis.

### Seminested Polymerase Chain Reaction (snPCR) Assay for Enterovirus VP1 gene

All primers were re-constituted in concentrations of 100μM and first round PCR was done in 50μL reactions. The first round PCR contained 0.5μL each of primers 224 and 222 (Nix et al., 2006), 10μL of Red load Taq, 10μL of cDNA and 29μL of RNase free water. A Veriti thermal cycler (Applied Biosystems, California, USA) was used for thermal cycling as follows; 94°C for 3 minutes followed by 45 cycles of 94°C for 30 seconds, 42°C for 30 seconds and 60°C for 60 seconds with ramp of 40% from 42°C to 60°C. This was then followed by 72°C for 7 minutes and held at 4°C till terminated.

The second round PCR was done in 30μL reactions. It contained 0.3μL each of forward and reverse primers (Figure 2), 6μL of Red load Taq, 5μL of first round PCR product and 18.4μL of RNase free water. As shown in the algorithm (Figure 1), six different second round PCR assays were done using the first round PCR product as template. Hence, all six second round PCR assays used different forward primers (Table 2) but the same reverse primer (AN88) (Figure 2). Based on the expected amplicon size (Figure 2), the PCR extension time for the second round assays were 30 seconds for primers AN89, 189 and 187 (WHO, 2015; Oberste et al., 2003; Nix et al., 2006) and 60 seconds for primers Sab-1, Sab-2 and Sab-3 (Sadeuh-Mba et al., 2013). The extension temperature was however retained at 60^0^C. Subsequently, PCR products were resolved on 2% agarose gel stained with ethidium bromide, and viewed using a UV transilluminator.

**Figure 2:**
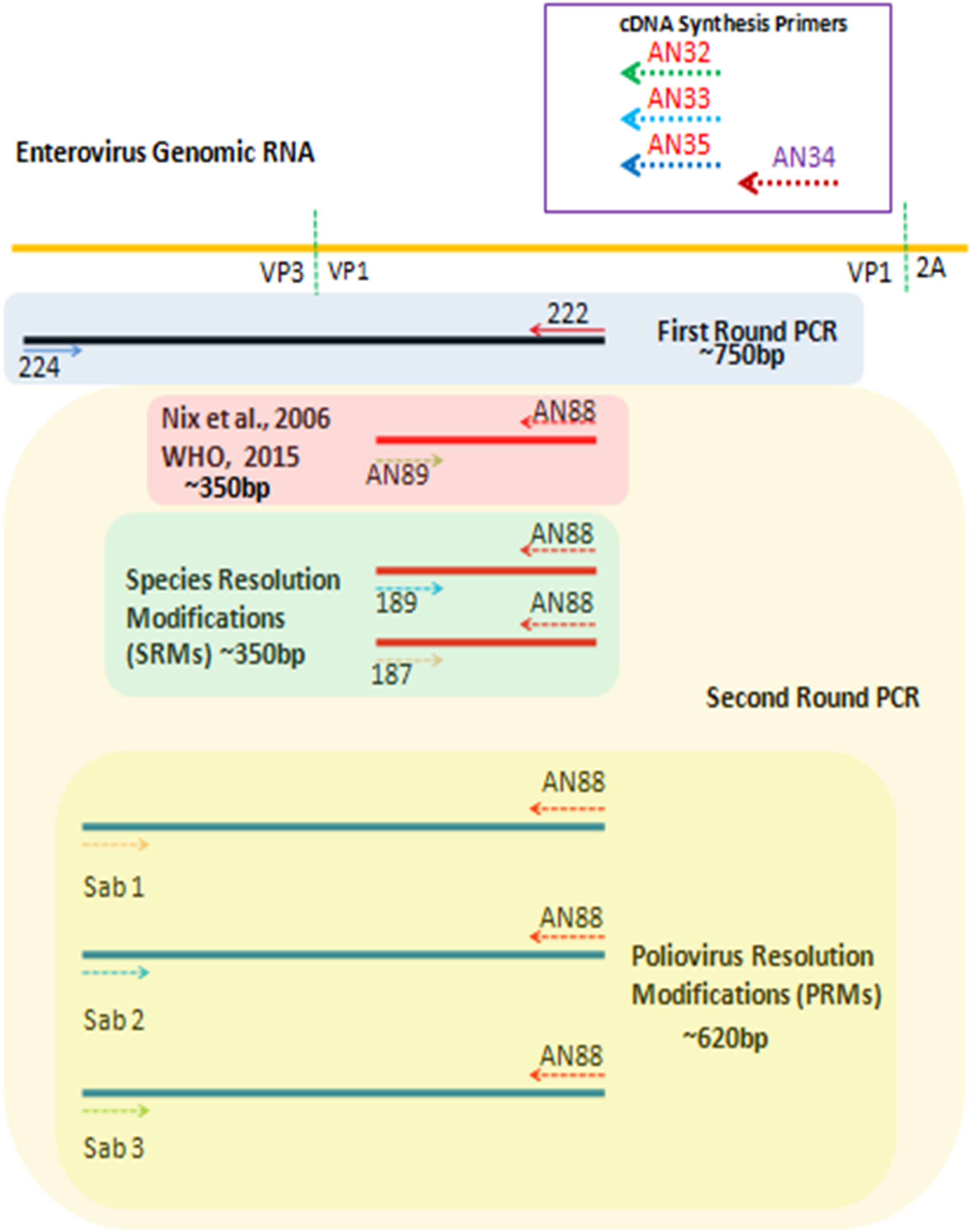
A schematic representation of the annealing sites of the different primers used in this study relative to the enterovirus genome and the consequent amplification product (arrows depict primers and their orientation).

**Table 2:**
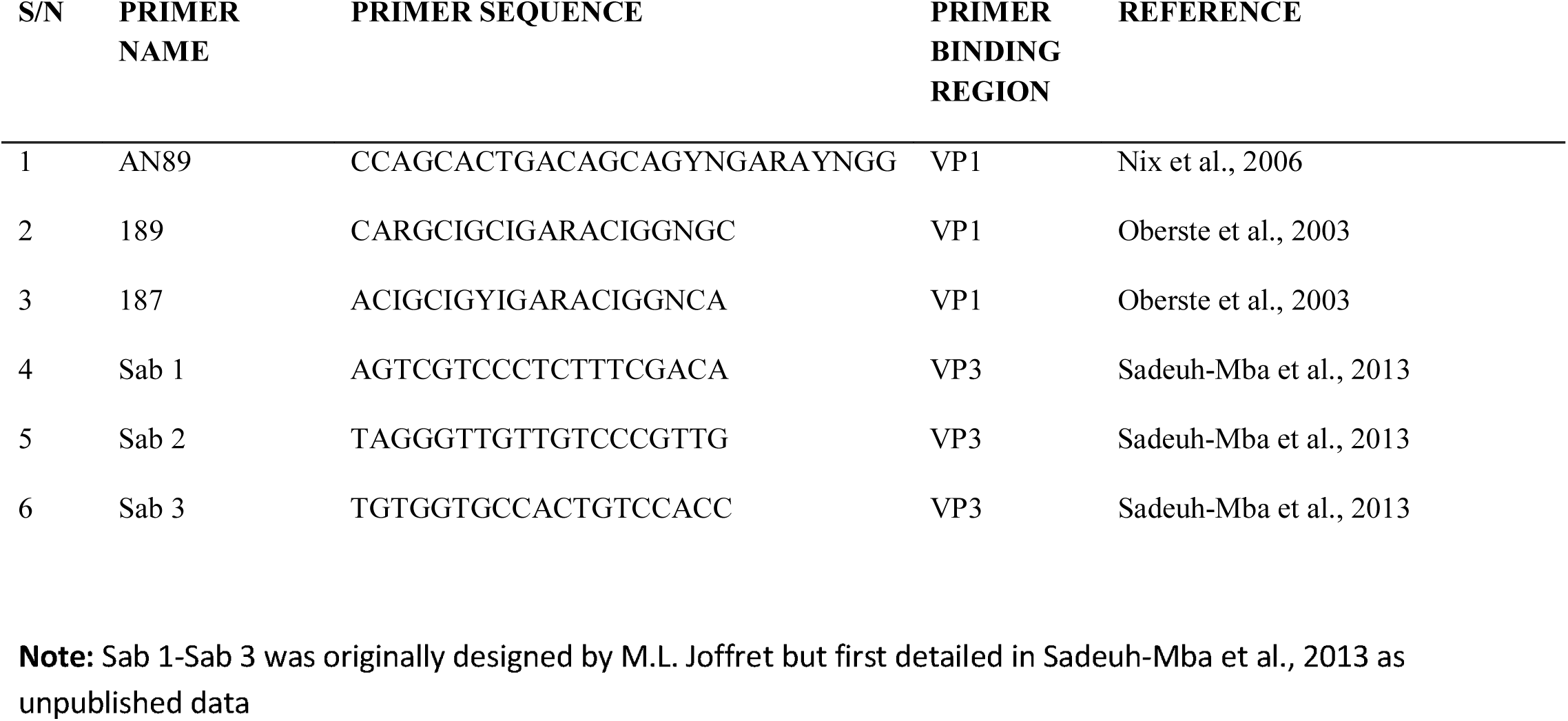
Sequences of the different forward primers used for second round PCR in this study.

### Nucleotide sequencing and enterovirus identification

All amplicons generated from the six second round PCR assays were shipped to Macrogen Inc, Seoul, South Korea, for purification and sequencing. The primers used for each of the second round PCR assays (Figure 2) were also used for sequencing, respectively. The identities of the sequenced isolates were determined using the enterovirus genotyping tool (Kroneman et al., 2011).

### Nucleic acid sequences described in this study

The accession numbers for the newly described sequence data generated in this study are **KX856914-KX856921**.

## RESULT

For isolates one to eight which were previously known to be EV-Bs, primer AN89, amplified only six of the eight samples (Table 1). It is noteworthy that the two isolates not amplified by primer AN89 were previously identified as E19. Also, the only previously identified E19 (sample 2) subsequently amplified by primer AN89 in this study was identified as E7 (Table 1). Further, the identity of the previously unidentified (sample 8) was shown to be E7, and all the other previously identified E7s and the E3 were confirmed (Table 1).

The Species Resolving Primers (SRPs [189 and 187[) identified the isolates to a large extent in accordance with their previously stated predilections (WHO, 2015). Hence, as expected primer 187 confirmed all but one (sample 4) of the eight (8) EV-B isolates as such. All its type identities were in accordance with that previously determined prior this study. It also identified the previously unidentified sample 8 as E7. Primer 189 however showed the presence of E20 in an isolate (sample 4) previously shown to be E19. The Poliovirus Resolving Primers (PRPs [Sab 1, 2 and 3]) however, did not amplify any of the isolates in samples 1 to 8 (Table 1).

For isolates nine to twelve which were all previously identified as CV-A13 (EV-C) (Faleye and Adeniji, 2015), Primer AN89 (Nix et al., 2006; WHO, 2015) detected and identified all as such. The SRPs also confirmed all as such. However, it is worth mentioning that primers 189 and 187 could not confirm samples 9 and 11 respectively because the sequence data were unexploitable. The PRPs did not amplify any of the isolates in samples 9 to 12 (Table 1).

For isolates 13 to 15 (previously identified as Sabin PV-1, 2 and 3 respectively), primer AN89 (Nix et al., 2006; WHO, 2015) detected all, and were subsequently confirmed as such. The SRPs also confirmed all as such. Similarly, primer 189 detected and identified all the three, but primer 187, though detected all three could only confirm sample 13 as PV-1. The remaining two were unexploitable due to bad sequence data. The PRPs also confirmed all as such. Precisely, the Sabin-1 primer only detected sample 13 and confirmed it to be PV-1. The Sabin-2 primer produced the expected amplicon size in both samples 13 and 14. The sequence data subsequently confirmed both to contain two different Sabin 2 viruses that are 99.65% similar (data not shown). In fact, the Sabin 2 in sample 13 has an Isoleusine (I) at position 143 of VP1 while that in sample 14 has an Asparagine (N) at the same position. The Sabin-3 primer also produced the expected amplicon size in both samples 13 and 15. The sequence data also confirmed both to contain two different Sabin-3 viruses that are 99.65% similar (data not shown). Hence, primers AN89 and the SRPs (189 and 187) did very well in identifying all three polioviruses. The PRPs (Sab1-Sab3) also confirmed the identity of the three samples. The PRPs however, further showed that sample 13, which was previously identified by the GPLN algorithm (Kilpatrick et al., 2009) as PV-1 and confirmed by primers AN89 and the SRPs as such, also contained PV-2 and PV-3 (Table 1).

For sample 16 which contained four EV-Cs (PV-1, PV-2, PV-3 and CV-A13), and two EV-Bs (E3 and E7), all the six primers produced their expected band sizes. Sequence data analysis however showed that while the amplicon from primer 189 produced unexploitable sequence data, primers AN89, 187, Sab-1, Sab-2 and Sab-3 detected PV-2, PV-1, PV-1, PV-2 and PV-3, respectively (Table 1). Otherwise stated, AN89 and the SRPs (primers 189 and 187) only detected PV-2 and PV-1 respectively from the mixture while the PRPs specifically detected all the three poliovirus serotypes. Importantly, the PV-2 and 3 detected in sample 16 were exactly those found in samples 14 and 15 respectively (Table 1).

## DISCUSSION

### Extending the Utility of the WHO recommended RT-snPCR assay

The results of this study confirm our previous findings (Faleye and Adeniji, 2015a, Faleye et al., 2016b) that though the Oberste et al., (2003) and the Nix et al., (2006) (recently recommended by the WHO [2015]) protocols are very sensitive for enterovirus detection and identification, both lack the capacity to resolve enterovirus co-infection. For example, samples 1-8 (Table 1) were all previously identified (Faleye and Adeniji, 2015b) using the Oberste et al., (2003) protocol. However, the results of this study showed that samples 2 and 4 were mixtures; a fact that was missed when the Oberste et al., (2003) protocol was used for identification (Faleye and Adeniji, 2015). Hence, in this study, the inclusion of the SRPs in the second round PCR of the WHO recommended RT-snPCR (WHO, 2015) assay enabled enterovirus co-infection detection and thereby improved the resolving capacity of the assay.

In the same light, by including poliovirus serotype specific forward primers (Sadeu-Mba et al., 2013) in the second round PCR of the WHO recommended RT-snPCR (WHO, 2015) assay, we were able to selectively and specifically detect and identify the different poliovirus serotypes. More importantly, this was accomplished in a situation where six different enterovirus types were present. The results of this study showed that in such instance, the RT-snPCR assay recommended by the WHO (WHO, 2015) detected only one of the six different enterovirus types present (Table 1; sample 16). It therefore confirmed that the WHO recommended RT-snPCR assay (WHO, 2015) might not be dependable for the resolution of enterovirus mixtures. Furthermore, the results of this study showed that the WHO recommended RT-snPCR assay (WHO, 2015) can be modified or tailored as described in this study (for detection and identification of the polioviruses) to specifically detect other enterovirus types especially in cases of co-infection. Such serotype-specific modifications would be very valuable for low-income economies as it will broaden the surveillance capacity of enterovirologists in such settings with minimal increase in cost.

### Sensitivity of the modified assay

It is pertinent to note that this modification appears to be more sensitive for poliovirus detection and identification than both the WHO RT-snPCR algorithm (Nix et al., 2006; WHO, 2015) and the current algorithm for poliovirus identification in use by the GPLN (Kilpatrick et al., 2009). For example, while the GPLN algorithm (Kilpatrick et al., 2009) and the WHO RT-snPCR algorithm (WHO, 2015) identified sample 13 as PV-1, the modification described here showed that sample 13 contained PV-1, 2 and 3. More importantly, sequence comparison showed that the PV-2 and 3 in sample 13, and those in samples 14 and 15 were different (data not shown). Consequent of this discovery, the initial result of this reference sample of environmental origin was retraced. It was then discovered that the results of the five L20B and one RD flask (WHO, 2004) showed that PV-1, 2 and 3 were all isolated from the parent environmental sample but in different flasks (unpublished data). However, the PV isolate in the L20B flask from which sample 13 was aliquoted was identified as PV-1 by the current GPLN poliovirus detection and identification algorithm (Kilpatrick et al., 2009). Altogether, this suggests that the isolate in sample 13, contained PV-1, 2 and 3 but PV-1 had a titre that is significantly higher than others and was consequently the type detected in sample 13 both by the GPLN algorithm and primers AN89, 189 and 187 (Table 1; sample 13). Further buttressing the influence of PV-1 titre hypothesis is the fact that the PV-2 and 3 in sample 13 could not be detected in the sample 16 mixture. Rather it was the PV-2 and 3 in samples 14 and 15 that were detected in sample 16 (Table 1).

Considering that about 25,000 genomic equivalents are required for the current GPLN algorithm to detect the presence of PV-1 (Arita et al., 2015), this finding is not unexpected. Rather it suggests that while the genomic equivalents of PV-1 in sample 13 might be up to the required, those of PV-2 and PV-3 are below and account for the inability of the assay to detect both. This has implications for the polio eradication and endgame strategic plan 2013-2018 (WHO, 2013) and particularly the WHO global action plan (GAP III) for poliovirus containment and sequential withdrawal of the Sabin strains (WHO, 2014). For example, in the course of Sabin PV-2 containment, isolates containing Sabin PV- 2 but with titre below the detection limit of the GPLN assay for Sabin PV-2 detection might be missed. It is therefore suggested that for containment, all isolates that contain any of the poliovirus types should be handled as potentially containing the other two types. Furthermore, to reduce the misclassification of mixed isolates with low titre components and consequently enhance the containment programme, effort should be put into increasing the sensitivity of the assays in use by the GPLN and others; like that described in this study. In addition, effort should also be put into mainstreaming serotype-independent NextGen sequencing strategies recently described for the polioviruses (Montmayeur et al., 2016) and other Species C members (Bessaud et al., 2016).

### Field implementation of the modified assay

It is imperative to mention that in this study; cell culture isolates were used and as such it might be difficult to remark on how the assay might fair for direct detection of polioviruses from clinical specimen. However, considering that the initial development of the RT-snPCR assay (Nix et al., 2006) recently recommended by the WHO (WHO, 2015) was done using isolates, we see no reason why implementing this algorithm with clinical specimen should be problematic. Furthermore, this assay does not in any way change the assay described by Nix and colleagues (Nix et al., 2006) and subsequently recommended by the WHO (WHO, 2015). Rather, considering its limitations with respect to resolving mixtures, this modification piggy-back on the first round PCR product of the Nix algorithm (Nix et al., 2006) in a bid to improve its mixture resolving capacity and thereby extend its utility.

In a polio-free world there might be reduced funding for poliovirus surveillance and only essential facilities (WHO, 2014) with appropriate safety mechanisms in place to avoid facility associated escape of poliovirus might be cleared for poliovirus isolation in cell culture. In such strictly regulated and potentially resource limited setting, the modification described in this study might be of significance because it allows the enlistment of nonessential facilities and others without the capacity or infrastructure for cell culture to participate in poliovirus surveillance. Particularly, in such settings, enterovirologists interested in nonpolio enteroviruses can continue their investigations, and also contribute significantly and specifically to poliovirus surveillance, by using the excess of their first round PCR product.

The modification described in this study is also valuable in regions where maintaining the reverse-cold chain for sample transport to the laboratory might constitute a bottle-neck. In such settings, using this modification as an addendum to the current cell culture based algorithm, even if only for cell culture negative samples, will improve the detection of poliovirus in samples that might no longer contain infectious particles on arrival in the laboratory. This might help reduce the incidence of orphan polioviruses by serving as an early warning system in determining regions where samples arrive in nonviable conditions. Such early detection can result in quick intervention and the consequent forestalling of the occurrence of orphan polioviruses. Also, like the RT-snPCR assay recently recommended by the WHO (2015), this algorithm will significantly reduce the turn-around time from sample arrival in the laboratory to availability of results to less than 48 hours. This is significantly less than the minimum of 10 days for the cell culture based GPLN algorithm (WHO, 2004) and is of immense value for early outbreak detection of a virus that only shows clinical manifestation in one out of 100 to 250 people infected (Nathanson and Kew, 2010).

### Limitations and Conclusion

The limit of detection of the modification described here is currently not known. Hence, effort is ongoing to conduct spiking experiments with plaque purified and titrated reference isolates in a bid to better define the sensitivity of the assay. Furthermore, it is crucial to mention that the sequence data this modification provides do not cover the entire VP1 region. As such, unlike the ECRA assay (Arita et al., 2015), the sequence data generated might not be sufficient for extensive molecular epidemiology. Consequently, this modification is currently proposed as addenda and not as substitute for either the current GPLN algorithm (WHO, 2004) or the other cell culture independent assays (Arita et al., 2015; Krasota et al., 2016) with the capacity to provide sequence data of the entire VP1 region. In addition, its value for early outbreak or extended vaccine virus circulation detection (currently in resource-limited settings and in a polio-free world) by increasing the number of laboratories that can contribute in a meaningful way to poliovirus surveillance is espoused.

## Acknowledgements

The authors would like to thank the members of staff of the WHO Environmental Surveillance Laboratory in Ibadan, Nigeria, for providing us with Sabin strains of poliovirus types 1 and 3.

## Compliance with ethical guidelines

Isolates recovered from environmental samples were analyzed in this study. Hence, this article does not contain any studies with human participants performed by any of the authors. The poliovirus 2 analysed in this study has been destroyed.

